# A new workflow combining R packages for statistical analysis of metabolites

**DOI:** 10.1101/848812

**Authors:** Paola G. Ferrario

## Abstract

In metabolomics, the investigation of an association between many metabolites and one trait (such as age in humans or cultivar in foods) is a central research question. On this topic, we present a complete statistical analysis, combining selected R packages in a new workflow, which we are sharing completely, according to modern standards and research reproducibility requirements. We demonstrate the workflow using a large-scale study with public data, available on repositories. Hence, the workflow can directly be re-used on quite different metabolomics data, when searching for association with one covariate of interest.

## 1 Introduction

The establishment of a relationship between the human metabolome and human basic traits (such as age or sex) is a key issue in human metabolomics. Accordingly, the analogous research question in food metabolomics concerns the relationship between food metabolome and food cultivar. See for instance [2, 7, 21, 20, 16, 6] and the articles cited there. However, the demonstration of such relationships is a challenging task, since the metabolomics data are complex and many issues are to be simultaneously managed, as we shall discuss later. In the literature of the previous decade, many attempts to gain information from metabolic data and systematically report results may be identified. However, as noted in [4], an excessive number of published studies are unclearly or incompletely reported, so that it is impossible to follow or replicate statistical analyses.

In this paper, we offer a reproducible workflow for analysis of metabolomics data in relation to one covariate of interest. The workflow is demonstrated by re-using well-documented data, to be found on the repository MetaboLights [9], as suggested in [4] as instrument to improve clarity and transparency in metabolomics analysis.

In order to gain an appreciation for the complexity of the data, note that the metabolites are scaled variously, reveal heterogeneous variances, may be correlated and may have ties, while some of them have values below a detection limit. Moreover, the data are detected on different platforms and they undergo platform-specific measurements and normalisations. Finally, in many studies, there are many more metabolites than probands, so that the so-called *p > N* problem arises (*p* number of metabolites, *N* sample size). On the other hand, one should consider that the covariate of interest can be modelled quantitatively as well as qualitatively, and, finally, a suitable model for the association between the metabolites and the covariate has to be set.

In the section Methods, the statistical procedures involved in the workflow will be explained in detail and the code of the workflow is presented. Finally, in the section Results, the output of the workflow is illustrated by applying it to large-scale data. The workflow is shared in form of reproducible code written in R and as knitr report, which can be found in the additional materials.

## 2 Methods

### 2.1 The model

Our aim is to detect the impact of a continuous covariate *X* (for example age) on *p* human metabolites *Y*_1_, *Y*_2_, …, *Y*_*p*_, where the problem dimension *p* is typically large in metabolomics and where the metabolites *Y*_*i*_ are sometimes left censored (because they have some values under a detection limit). The association between a covariate of interest *X* and the *p* metabolites *Y*_*i*_ for *i* = 1, …, *p* can be expressed by the equations

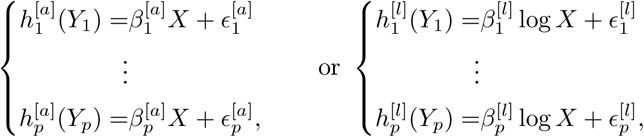

where *h*_*i*_ are different, strictly monotonically increasing transformations of *Y*_*i*_, and *β*_*i*_ are effects, whose interpretation arises from the *a priori* chosen error distributions of *ϵ*_*i*_. Moreover, as indicated by [19], the covariate *X* is modified by its logarithms, yielding in total two regression models per metabolite *Y*_*i*_, respectively, expressed by notation [a] (for ari, arithmetic) and [l] (for log, logarithmic for *X >* 0), respectively. The basic idea is again to be receptive for different association patterns between the metabolites and the covariate of interest as in [19]. It should be noted that beside the logarithm other modifications (or so called metameters) of the covariate can also be considered. After these steps, our research question will be approached by estimating the function *h*_*i*_ from the data. This estimation is embedded in a maximum likelihood framework and can be facilitated by parametrization of *h*, for our aims in terms of its basis functions by Bernstein polynomials, for instance. The conditional distribution of *Y* given *X* is estimated from a flexible parametric model using Bernstein polynomials. The resulting estimate of *h* is called *most likely transformation*. For a deeper discussion of this approach, we refer the reader to [11].

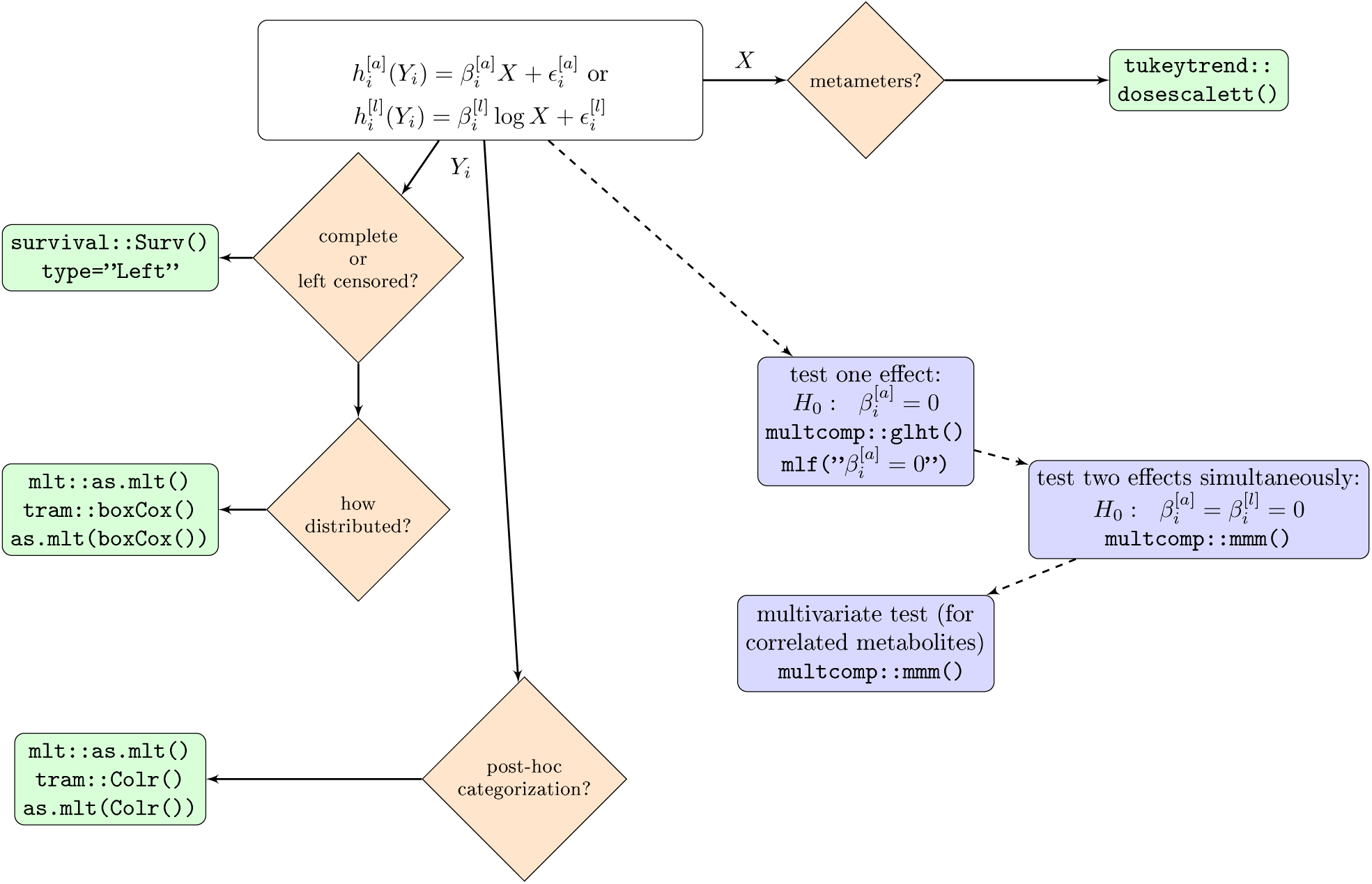

The null hypothesis is that there is no association between *Y*_*i*_ and *X* or between *Y*_*i*_ and log(*X*). The alternative hypothesis is that at least one model shows association, that is:

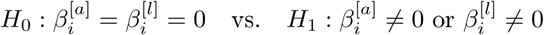

The equivalent multivariate hypothesis generalizes this for all metabolites *Y*_*i*_, *i* = 1, …, *p*. For the in total 2 ∗ *p* models, the correlation between the marginal test statistics is calculated by multiple marginal models. For a deeper discussion of this approach, we refer the reader to [14]. The corresponding test statistics are i.i.d.; many estimation techniques can be adopted from the theory on maximum likelihood estimation. Having set the seed, the quantiles are calculated using some numerical approximations. We refer the reader to [10] for further details, together with the associated references cited.

At this point we notice that also an additional approach could be considered if one is interested in any categorisazion of the metabolites. The approach is the so-called continuous outcome logistic regression, a technique that is recently proposed by [13] and represents a valid alternative to post-hoc categorizations (e.g. the use of four dietary reference quartiles [5]) that is still widely used despite all warnings [1, 15]. This type of regression makes it possible to consider the association between the covariate of interest and the metabolites by odds ratios that can be evaluated for all potential values or cut-off of the metabolites.

For this setting, the workflow presented involves the R packages `survival, tram, mlt, multcomp`. The flowchart in the full-page-figure schematically displays the main principles behind the workflow together with the packages. Orange diamonds represent mandatory questions regarding data characteristics or analysis strategy. Green rectangles list the R packages and functions involved as proposed solution. Blue rectangles between dashed lines indicate possible inference steps.

### 2.2 Data example

In order to explain the code of the workflow step-by-step, we use a small-scale example consisting of three (real) variables picked up for our demonstration from the KarMeN metabolomics and nutrition cross-sectional study, published by [3, 7]. The three (real) variables are: the metabolite GUDCA (Glycoursodeoxycholic Acid, bile acid measured on LC-MS in human plasma), the metabolite C10.0 (Decanoic acid, fatty acid measured on GC-MS in human plasma) and the age of 291 probands. The aim is to investigate association between age and the metabolites GUDCA and C10.0, respectively and jointly. The results of this analysis will be given in form of a table containing the estimated effects and the confidence intervals quantifying association (compare Table 1). For a minimal statistical description of the three variables involved, Figure 1 displays their empirical distribution functions (estimated by Kaplan-Meier estimator in case of censoring). It should be noted that only for the sake of code explanation we chose these two metabolites; the reality is much more complex: in fact, the KarMeN data set consists of more than 1000 metabolites, measured on many platforms and following different quality control protocols.

**Table 1:** Simultaneous confidence intervals.

| metabolite | metameter | effect | lower limit | upper limit |
| --- | --- | --- | --- | --- |
| GUDCA | arithmetic | -0.017 | -0.025 | -0.009 |
| GUDCA | logarithmic | -0.757 | -1.102 | -0.413 |
| C10.0 | arithmetic | 0.003 | -0.005 | 0.011 |
| C10.0 | logarithmic | 0.121 | -0.215 | 0.457 |

**Figure 1:**
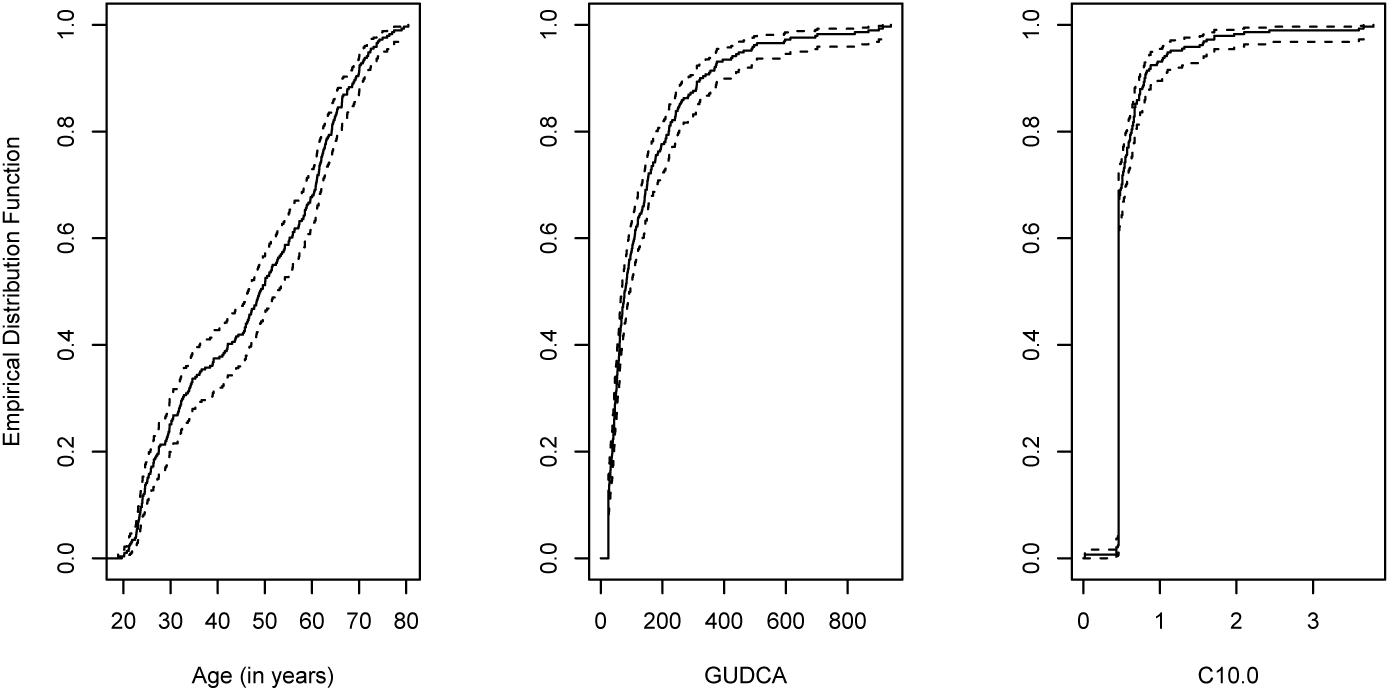
Empirical distribution functions (estimated by Kaplan-Meier estimator in case of censoring) for age, GUDCA, and C10.0 with dotted lines for the two-sided 0.95 confidence intervals

## 2.3 Code explanation

In the first code lines, we involve the two metabolites considering their specific properties. The measurement of GUDCA is affected by a limit of quantification (LOQ) of 25: this is considered modelling GUDCA as left censored. The measurement of C10.0 is affected by many repeated values, so-called ties. Ties are generally an issue; many methods exist for dealing with them; compare for instance [12]. In many cases ties result from insufficient precision in measurement, in other cases ties comes from the discrete character of a variable. One method to break ties is to add a uniformly distributed random variable. Another way to break them is to regard them as censored, interval variables. The R function `survival::Surv` can manage this.

~~~
library(“survival”)
### (I) metabolite GUDCA with left censoring at value 25
dd$GUDCAsurv <- with(dd, Surv(GUDCA, event = GUDCA > 25,
     type = “left”))
### (II) metabolite C10.0 with lots of ties
dd$C10.0surv <- with(dd, Surv(time = C10.0 - 0.005,
     time2 = C10.0 + 0.005,
     type = “interval2”))
~~~

As indicated by Tukey [19], we consider not only the age of the probands but also its logarithm: This means that we have two association models, one with age as covariate of interest *X* =age, the other with its logarithm log(*X*) = log(age). The metameters can simply be calculated directly but also by the R function `tukeytrend::dosescalett`, as demonstrated here

~~~
## age metameters
TukeyMetam <- dosescalett(data = dd, dose = “Age”,
     scaling = c(“ari”, “log”))
m_dat <-TukeyMetam$data # Tukeys dose metameters
~~~

In the next chunk, the support of the variable GUDCA is considered, taking into account the 95% quantile of the variable. This support is involved in the definition of the domain for the Bernstein polynomials for calculation of the most likely transformation

~~~
 ## support of baseline transformation
suGUDCA <- quantile(survfit(GUDCAsurv ∼ 1, data = dd),
     prob = c(.01, .95))$quantile
lfma <- as.mlt(BoxCox(GUDCAsurv ∼ Ageari, data = m_dat,
     support = suGUDCA)
lfmA <- as.mlt(BoxCox(GUDCAsurv ∼ Agelog, data = m_dat,
     support = suGUDCA)
~~~

The association between the metabolite GUDCA and age in its two metameters is investigated by performing the following test: the null hypothesis postulates that age and the metabolite have no association (`ari = “Ageari=0”`) or that log(age) and the metabolite have no association ((`log = “Agelog=0”`). As a result either a bivariate test statistic or two marginal test statistics can be considered per metabolite. The correlation between the two marginal test statistics is calculated (by multiple marginal models, R function (`multcomp::mmm`) and taken into account for the joint inference (by general linear hypothesis, R function `multcomp::glht`)

~~~
## joint inference for all two metameters
maxT <- glht(mmm(ari = fma, log = fmA),
     mlf(ari = “Ageari=0”, log = “Agelog=0”))
~~~

The same procedures are repeated for metabolite C10.0. Finally, by `multcomp::mmm` one can consider a multivariate hypothesis test, where the association between both metabolites in the two metameters and *X* is investigated simultaneously. Finally, the confidence intervals are calculated.

~~~
####### bivariate multiple metabolites test
maxTbiv <-glht(mmm(ari=fma, log=fmA, ari2=fma2, log2=fmA2),
     mlf(ari=“Ageari=0”, log=“Agelog=0”,
      ari2=“Ageari=0”, log2=“Agelog=0”))
~~~

The reader can find the code in the additional files.

## Results

In order to illustrate the results of our workflow, we revisited the data presented in the article [18], which are available online in the data repository MetaboLights [9] and on Workflow4Metabolomics.org [8] (here one list of recommended data repositories for metabolomics https://www.nature.com/sdata/policies/repositories).

Specifically, this large scale-study investigates the impact of age as covariate of interest on the urinary metabolome, described by 120 metabolites measured by LC-HRMS. The cohort consisted of 183 adults in a cross-sectional design. The impact of age is considered simultaneously on all the metabolites, i.e., in a multivariate approach. The complete results of applying the workflow to the data can be visualized with one plot: Figure 2 draws one confidence interval for each metabolite; together, the confidence intervals for all 120 metabolites are ordered by increasing effects for linear association with age, allowing scale-independent interpretation. The confidence intervals transgressing the vertical line (for effect=0) describe the metabolites associated with age; specifically, they show a positive correlation when they are at the right-hand side of the vertical line and a negative correlation when they are on the left. Moreover, the effects are listed in table (to be found in the supplementary material).

**Figure 2:**
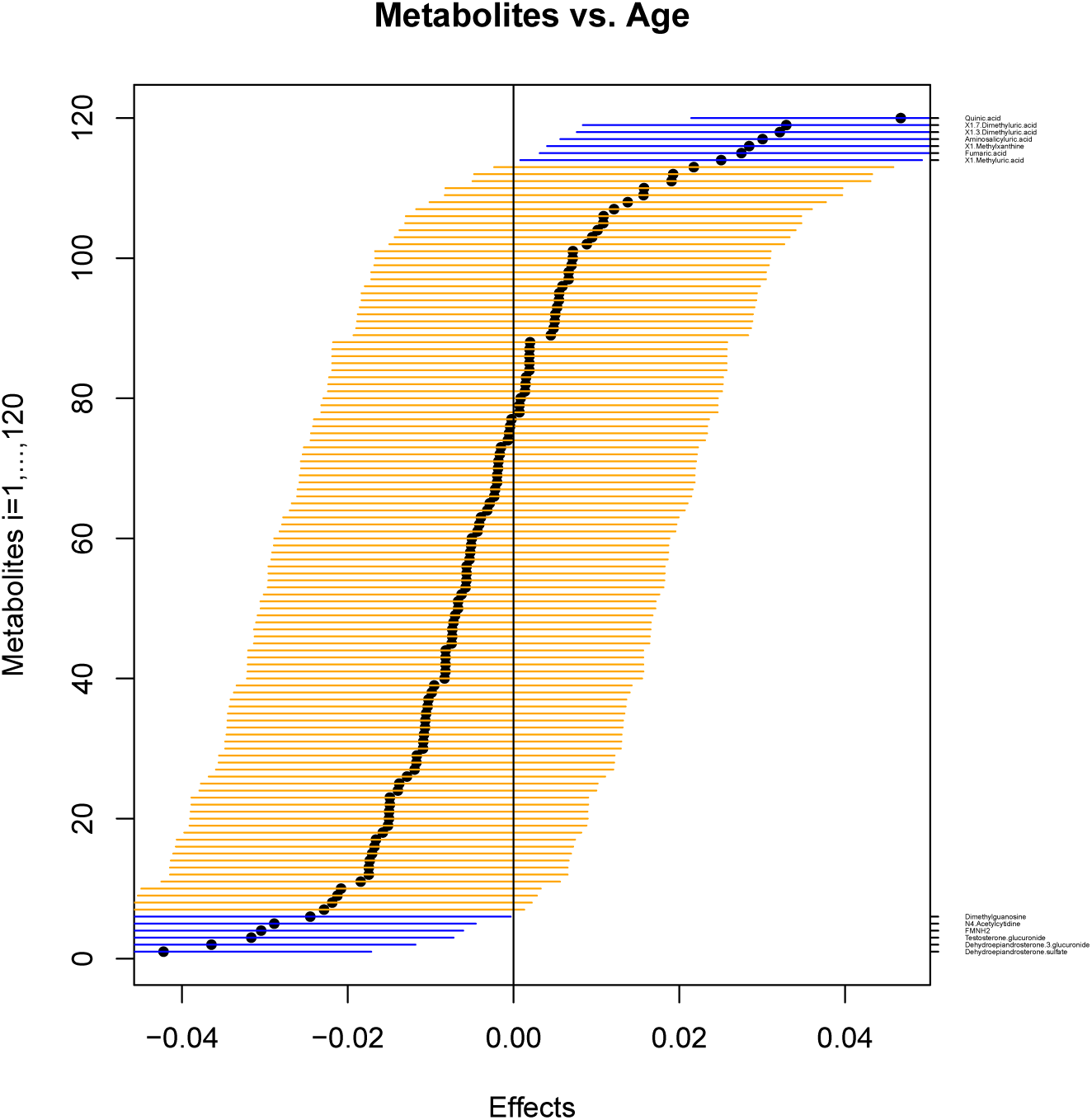
Impact of age on 120 urine metabolites for 183 adults under a linear model, visualized by confidence intervals, ordered by increasing effects, given as horizontal lines around black circles. The blue lines are the metabolites that show association with age.

## Discussion

The workflow has the following general attributes:

- There are no a priori assumptions about the distribution of *Y*_*i*_, since it is unrealistic to assume the same error distribution for each of the *p* metabolites. Instead, a metabolite-specific data-driven transformation function of the outcome will be involved by mlt.
- mlt provides a comparable analysis of such differently scaled metabolites and therefore enables scale-independent interpretation of the results, which is particularly helpful when searching for association between one specific covariate of interest and the multiple metabolites simultaneously.
- The workflow takes into account the fact that the *p* metabolites are a mixture of completely measured metabolites but also left censored metabolites (with values below the limit of detection or quantification), which is an intrinsic property of the chemical measurement of metabolites. Moreover, some metabolites have many ties, and these too are integrated in analysis.
- Different association patterns between the metabolites *Y*_*i*_ and the covariate of interest *X* are “allowed”: The workflow is also able to detect non-linear associations.
- Each metabolite gets an own “suitable” model for association with the covariate of interest as result of a maximization process without need for explicit formulation of some parameters, so that the model can be said to be found in an unsupervised way.
- The adjustment for multiple comparisons considers another intrinsic property of the metabolites, namely that they are often correlated in subgroups. This information has been included and leads to adjustments for multiplicity that can be less conservative compared to approaches that ignore it.

## Conclusion

We have proposed a workflow for statistical analysis of metabolites in relation to one covariate of interest. According to the recommendations of Open Science [17], we are sharing the code in its entirety. We have demonstrated the workflow using public data, available on recognized repositories.

The workflow is based on combinations of many R packages - `survival, tram, mlt, multcomp`, considering the fact that the metabolomics data are differently scaled, platform-dependent, heterogeneous, and sometimes not detectable under a certain limit. Considering the most likely transformation function for each metabolite, we enable the metabolites to be differently distributed, left censored and have different patterns of association with the covariate of interest, or with its logarithm, too. According to modern statistical approaches, we have included the correlation between metabolites in order to be less conservative than by classical approaches (like Bonferroni-correction) when adjusting for multiplicity. A possible limitation of the workflow occurs if there is a large proportion of values under a detection limit. In this case it could be difficult to find the Bernstein polynomials and hence to calculate the most likely transformation. With the continuous outcome logistic regression, we are suggesting a possible generalization of the procedure, should there be interest in categorizing the metabolites.

We have shared the code for a presented example and for analysis of large-scale data: anybody is able to re-use the workflow on the proposed data and on other data, thus enabling research reproducibility, transparency and clarity and subsequently progress in metabolomics research.

## Supporting information

Additional file 3: Pdf file based on the Rnw-Code in additional file 2.

Additional file 2: Rnw-Code that demonstrates the application of the workflow on a large-scale study (impact of age on the urinary metabolome).

Additional file 1: R-Code that demonstrates the workflow using the R packages survival, tram, mlt and multcomp, by means of the example data.

## Additional files

1. Additional file 1: R-Code that demonstrates the workflow using the R packages `survival`, `tram`, `mlt` and `multcomp`, by means of the example data.
2. Additional file 2: Rnw-Code that demonstrates the application of the workflow on a large-scale study (impact of age on the urinary metabolome).
3. Additional file 3: Pdf file based on the Rnw-Code in additional file 3.

## Acknowledgements

I gratefully acknowledge Ludwig Hothorn for many helpful suggestions and for his active interest in the publication of this paper. This paper is based on a co-work with Ludwig Hothorn as part of a consultant agreement between him and the Max Rubner-Institut (with Internal Report, October 2017).

I wish to express my special thanks to Torsten Hothorn for his refinements in the R-code that made it clearer and more elegant.

