## Additional file 3: Pdf file based on the Rnw-Code in additional file 2. for "A new workflow combining R packages for statistical analysis of metabolites"

### Application of the workflow to a real, available dataset: Urinary metabolome variation with age

#### Contents

**1 Aim** 1

**2 Data** 1

**3 A little descriptive statistics** 1

**4 Re-analysation of data** 3

#### 1 Aim

The aim of this report is to re-analyse large-scale data that are available in literature using the presented workflow. Thus, we demonstrate how the workflow works in practice, which assumptions are required and how the results can be represented in form of, for instance, confidence intervals, plots, lists of p-values, etc....The code is completely reproducible.
To this end, we considered a cohort consisting of 120 analytes measured by LC-HRMS for 183 adults (compare the section below). In particular, we focussed on the association between the analytes and age as continuous covariate of interest.

#### 2 Data

Consider the article:
Thévenot et al. (2015) Analysis of the Human Adult Urinary Metabolome Variations with age, Body Mass Index, and Gender by Implementing a Comprehensive Workflow for Univariate and OPLS Statistical Analyses. *J Proteome Res*, **14(8)**, 3322-3335.
DOI: `10.1021/acs.jproteome.5b00354`
`https://pubs.acs.org/doi/10.1021/acs.jproteome.5b00354` and the data available together with the article

- `sacurine-neg_sampleMetadata.csv`, downloaded from `https://workflow4metabolomics.org/dataset_sacurine` (October 31th, 2019)
- `m_sacurine_v2_maf.csv`, downloaded from `https://www.ebi.ac.uk/metabolights/MTBLS404` (October 31th, 2019)

#### 3 A little descriptive statistics

Draw gender-specific age and BMI distributions

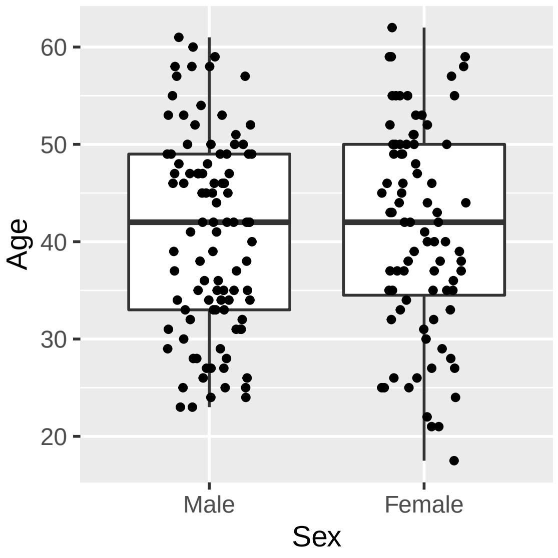


Figure 1: Gender-specific age distribution.

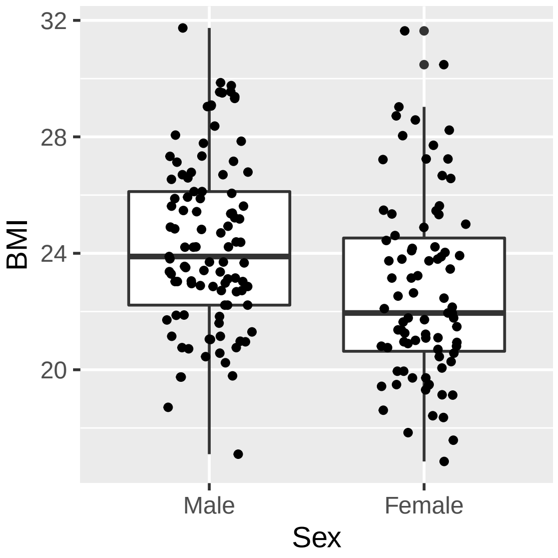


Figure 2: Gender-specific BMI distribution.

### 4 Re-analysation of data

Is there any association between each metabolite and age or between each metabolite and log(age), respectively?
Run for this the workflow on the data

```
analyt <- make.names(analyt_orig <- colnames(data.thevenot)[-(1:4)])
colnames(data.thevenot)[-(1:4)] <- analyt

dd <- dosescalett(data = data.thevenot, dose = "age",
                  scaling = c("ari","log"))$data

make_str <- function(a) {
  cm_ari <- as.mlt(BoxCox(as.formula(paste(a, " ~ ageari")), data = dd))
  cm_log <- as.mlt(BoxCox(as.formula(paste(a, " ~ agelog")), data = dd))
  cm_mmm <- glht(mmm(ari = cm_ari, log = cm_log),
                 mlf(ari = "ageari=0", log = "agelog=0"))
  list(cf = confint(cm_mmm)$confint,
       pvals = summary(cm_mmm)$test$pvalue,
       logLik = c(ageari = logLik(cm_ari),
                  agelog = logLik(cm_log)))
}
res <- lapply(analyt, make_str)
```

| Analyte | Age | Effect | Confint | p-Value | logLik |
|---|---|---|---|---|---|
| Quinic acid | ari | 0.05 | ( 0.03– 0.06) | <0.001 | -3055.40 |
| | log | 1.79 | ( 1.20– 2.38) | <0.001 | -3055.31 |
| Dehydroepiandrosterone sulfate | ari | -0.04 | (-0.06—0.03) | <0.001 | -3130.04 |
| | log | -1.60 | (-2.19—1.02) | <0.001 | -3130.23 |
| Dehydroepiandrosterone 3-glucuronide | ari | -0.04 | (-0.05—0.02) | <0.001 | -2473.95 |
| | log | -1.31 | (-1.88—0.74) | <0.001 | -2475.43 |
| 1,3-Dimethyluric acid | ari | 0.03 | ( 0.02– 0.05) | <0.001 | -2978.91 |
| | log | 1.30 | ( 0.73– 1.87) | <0.001 | -2977.89 |
| 1,7-Dimethyluric acid | ari | 0.03 | ( 0.02– 0.05) | <0.001 | -3162.10 |
| | log | 1.29 | ( 0.72– 1.87) | <0.001 | -3161.58 |
| Testosterone glucuronide | ari | -0.03 | (-0.05—0.02) | <0.001 | -2426.60 |
| | log | -1.16 | (-1.73—0.59) | <0.001 | -2427.53 |
| Aminosalicyluric acid | ari | 0.03 | ( 0.02– 0.04) | <0.001 | -2735.07 |
| | log | 1.17 | ( 0.60– 1.74) | <0.001 | -2734.57 |
| FMNH2 | ari | -0.03 | (-0.05—0.02) | <0.001 | -2049.05 |
| | log | -1.10 | (-1.66—0.53) | <0.001 | -2050.00 |
| 1-Methylxanthine | ari | 0.03 | ( 0.01– 0.04) | <0.001 | -3176.16 |
| | log | 1.16 | ( 0.59– 1.73) | <0.001 | -3175.46 |
| N4-Acetylcytidine | ari | -0.03 | (-0.04—0.01) | <0.001 | -2515.49 |
| | log | -1.07 | (-1.64—0.51) | <0.001 | -2515.94 |
| Fumaric acid | ari | 0.03 | ( 0.01– 0.04) | <0.001 | -2515.98 |
| | log | 1.01 | ( 0.44– 1.57) | <0.001 | -2516.51 |
| 1-Methyluric acid | ari | 0.03 | ( 0.01– 0.04) | <0.001 | -3241.22 |
| | log | 1.01 | ( 0.45– 1.57) | <0.001 | -3240.60 |

| | | | | | |
|---|---|---|---|---|---|
| Dimethylguanosine | ari | -0.02 | (-0.04—0.01) | <0.001 | -2652.98 |
| | log | -0.90 | (-1.46—0.33) | 0.0016 | -2653.48 |
| 6-(carboxymethoxy)-hexanoic acid | ari | -0.02 | (-0.04—0.01) | 0.0020 | -2662.46 |
| | log | -0.85 | (-1.41—0.29) | 0.0026 | -2662.90 |
| Methylinosine | ari | -0.02 | (-0.04—0.01) | 0.0030 | -2308.39 |
| | log | -0.79 | (-1.34—0.23) | 0.0054 | -2308.92 |
| Pyrroledicarboxylic acid | ari | 0.02 | ( 0.01– 0.04) | 0.0032 | -2496.21 |
| | log | 0.82 | ( 0.26– 1.38) | 0.0036 | -2496.33 |
| N-Acetyltryptophan isomer 3 | ari | -0.02 | (-0.04—0.01) | 0.0040 | -2721.70 |
| | log | -0.80 | (-1.36—0.24) | 0.0045 | -2721.85 |
| Aspartic acid | ari | -0.02 | (-0.04—0.01) | 0.0049 | -2741.63 |
| | log | -0.74 | (-1.30—0.18) | 0.0086 | -2742.15 |
| Pentose | ari | 0.02 | ( 0.00– 0.03) | 0.0089 | -3131.41 |
| | log | 0.73 | ( 0.17– 1.28) | 0.0100 | -3131.51 |
| Acetylphenylalanine | ari | 0.02 | ( 0.00– 0.03) | 0.0096 | -2616.93 |
| | log | 0.73 | ( 0.17– 1.28) | 0.0100 | -2616.92 |
| 4-Hydroxybenzoic acid | ari | -0.02 | (-0.03—0.00) | 0.012 | -2419.62 |
| | log | -0.67 | (-1.22—0.11) | 0.018 | -2419.83 |
| gamma-Glu-Leu | ari | -0.02 | (-0.03—0.00) | 0.018 | -2553.13 |
| | log | -0.62 | (-1.18—0.07) | 0.028 | -2553.42 |
| gamma-Glu-Ile | ari | -0.02 | (-0.03—0.00) | 0.018 | -2553.13 |
| | log | -0.62 | (-1.18—0.07) | 0.028 | -2553.42 |
| 3-Methylcrotonylglycine | ari | -0.02 | (-0.03—0.00) | 0.019 | -2903.84 |
| | log | -0.64 | (-1.20—0.08) | 0.023 | -2904.03 |
| N-Acetyltryptophan | ari | 0.02 | ( 0.00– 0.03) | 0.034 | -2334.38 |
| | log | 0.66 | ( 0.10– 1.21) | 0.020 | -2333.92 |
| Pantothenic acid | ari | -0.02 | (-0.03—0.00) | 0.025 | -2903.71 |
| | log | -0.65 | (-1.21—0.10) | 0.021 | -2903.53 |
| Pyridoxic acid isomer 1 | ari | -0.02 | (-0.03—0.00) | 0.021 | -3094.71 |
| | log | -0.63 | (-1.19—0.08) | 0.025 | -3094.70 |
| N-Acetyl-aspartic acid | ari | -0.02 | (-0.03—0.00) | 0.023 | -2437.82 |
| | log | -0.64 | (-1.19—0.08) | 0.024 | -2437.88 |
| Threonic acid | ari | -0.01 | (-0.03—0.00) | 0.043 | -3182.11 |
| | log | -0.61 | (-1.17—0.05) | 0.031 | -3181.78 |
| Erythronic acid | ari | -0.01 | (-0.03—0.00) | 0.043 | -3182.11 |
| | log | -0.61 | (-1.17—0.05) | 0.031 | -3181.78 |
| Methoxysalicylic acid isomer | ari | 0.02 | ( 0.00– 0.03) | 0.033 | -2524.35 |
| | log | 0.60 | ( 0.05– 1.16) | 0.033 | -2524.09 |
| Pyridylacetylglycine | ari | -0.02 | (-0.03—0.00) | 0.033 | -2744.36 |
| | log | -0.60 | (-1.15—0.04) | 0.035 | -2744.41 |
| Heptylmalonic acid | ari | -0.02 | (-0.03—0.00) | 0.04 | -2932.10 |
| | log | -0.56 | (-1.11—0.00) | 0.05 | -2932.28 |
| 5-Hydroxyindoleacetic acid | ari | -0.02 | (-0.03—0.00) | 0.043 | -2411.83 |
| | log | -0.54 | (-1.09– 0.02) | 0.059 | -2412.11 |
| Hydroxysuberic acid isomer 2 | ari | -0.02 | (-0.03—0.00) | 0.043 | -2460.11 |
| | log | -0.57 | (-1.13—0.01) | 0.044 | -2460.30 |
| Hydroxysuberic acid isomer 1 | ari | -0.01 | (-0.03– 0.00) | 0.063 | -2483.96 |
| | log | -0.57 | (-1.12—0.01) | 0.045 | -2483.68 |
| o-Hydroxyphenylacetic acid | ari | -0.01 | (-0.03– 0.00) | 0.060 | -2712.69 |
| | log | -0.55 | (-1.10– 0.01) | 0.054 | -2712.60 |

| | | | | | |
|---|---|---|---|---|---|
| Threo-3-Phenylserine | ari | 0.01 | (-0.00– 0.03) | 0.063 | -2922.50 |
| | log | 0.53 | (-0.02– 1.09) | 0.059 | -2922.43 |
| Glycocholic acid isomer 1 | ari | -0.01 | (-0.03– 0.00) | 0.082 | -1985.71 |
| | log | -0.41 | (-0.96– 0.14) | 0.151 | -1986.18 |
| Deoxyhexose | ari | 0.01 | (-0.00– 0.03) | 0.10 | -2558.72 |
| | log | 0.46 | (-0.10– 1.01) | 0.11 | -2558.76 |
| Tryptophan | ari | -0.01 | (-0.03– 0.00) | 0.11 | -2528.43 |
| | log | -0.42 | (-0.97– 0.14) | 0.15 | -2528.66 |
| 2-Hydroxybenzyl alcohol | ari | 0.01 | (-0.00– 0.03) | 0.15 | -2356.93 |
| | log | 0.45 | (-0.10– 1.01) | 0.11 | -2356.77 |
| 3,7-Dimethyluric acid | ari | -0.01 | (-0.03– 0.00) | 0.11 | -2715.96 |
| | log | -0.39 | (-0.94– 0.16) | 0.17 | -2716.33 |
| Kynurenic acid | ari | -0.01 | (-0.03– 0.00) | 0.12 | -2934.42 |
| | log | -0.41 | (-0.96– 0.14) | 0.15 | -2934.61 |
| 2-Aminoadipic acid | ari | -0.01 | (-0.03– 0.00) | 0.16 | -2336.25 |
| | log | -0.43 | (-0.98– 0.13) | 0.14 | -2336.19 |
| 2,2-Dimethylglutaric acid | ari | -0.01 | (-0.03– 0.00) | 0.14 | -2581.16 |
| | log | -0.38 | (-0.93– 0.17) | 0.18 | -2581.33 |
| p-Hydroxymandelic acid | ari | -0.01 | (-0.03– 0.00) | 0.14 | -2764.68 |
| | log | -0.39 | (-0.94– 0.17) | 0.18 | -2764.83 |
| Hydroxyphenyllactic acid | ari | 0.01 | (-0.00– 0.03) | 0.15 | -3192.65 |
| | log | 0.40 | (-0.15– 0.95) | 0.16 | -3192.79 |
| Acetaminophen glucuronide | ari | -0.01 | (-0.03– 0.00) | 0.15 | -2628.29 |
| | log | -0.10 | (-0.66– 0.45) | 0.76 | -2628.34 |
| Oxoglutaric acid | ari | -0.01 | (-0.03– 0.00) | 0.15 | -3027.88 |
| | log | -0.35 | (-0.90– 0.20) | 0.22 | -3028.10 |
| N2-Acetylaminoadipic acid | ari | -0.01 | (-0.03– 0.00) | 0.15 | -2686.55 |
| | log | -0.39 | (-0.94– 0.16) | 0.17 | -2686.61 |
| 5-Sulfosalicylic acid | ari | 0.01 | (-0.00– 0.02) | 0.20 | -2807.74 |
| | log | 0.41 | (-0.15– 0.96) | 0.15 | -2807.52 |
| Asp-Leu/Ile isomer 2 | ari | -0.01 | (-0.02– 0.00) | 0.16 | -2467.47 |
| | log | -0.33 | (-0.88– 0.22) | 0.24 | -2467.78 |
| 3-Indole carboxylic acid glucuronide | ari | -0.01 | (-0.02– 0.00) | 0.17 | -2842.79 |
| | log | -0.38 | (-0.93– 0.17) | 0.18 | -2842.87 |
| Valerylglycine isomer 1 | ari | -0.01 | (-0.02– 0.00) | 0.19 | -2638.99 |
| | log | -0.39 | (-0.94– 0.16) | 0.17 | -2638.94 |
| Benzoic acid isomer | ari | 0.01 | (-0.00– 0.02) | 0.18 | -2682.48 |
| | log | 0.37 | (-0.19– 0.92) | 0.20 | -2682.61 |
| 4-Acetamidobutanoic acid isomer 2 | ari | -0.01 | (-0.02– 0.00) | 0.20 | -3120.34 |
| | log | -0.36 | (-0.91– 0.19) | 0.21 | -3120.37 |
| Hippuric acid | ari | 0.01 | (-0.01– 0.02) | 0.24 | -3575.89 |
| | log | 0.35 | (-0.20– 0.91) | 0.22 | -3575.82 |
| Gluconic acid and/or isomers | ari | -0.01 | (-0.02– 0.01) | 0.28 | -2932.97 |
| | log | -0.33 | (-0.88– 0.22) | 0.25 | -2932.89 |
| 4-Acetamidobutanoic acid isomer 3 | ari | -0.01 | (-0.02– 0.01) | 0.28 | -2867.23 |
| | log | -0.33 | (-0.88– 0.23) | 0.26 | -2867.18 |
| Glu-Val | ari | -0.01 | (-0.02– 0.01) | 0.27 | -2715.28 |
| | log | -0.32 | (-0.87– 0.23) | 0.27 | -2715.28 |
| Sebacic acid | ari | -0.01 | (-0.02– 0.01) | 0.27 | -2940.18 |
| | log | -0.26 | (-0.81– 0.29) | 0.36 | -2940.37 |

| | | | | | |
|---|---|---|---|---|---|
| Glycocholic acid isomer 3 | ari | -0.01 | (-0.02– 0.01) | 0.28 | -2161.06 |
| | log | -0.24 | (-0.79– 0.31) | 0.42 | -2161.33 |
| p-Anisic acid | ari | -0.01 | (-0.02– 0.01) | 0.38 | -2865.82 |
| | log | -0.30 | (-0.85– 0.25) | 0.30 | -2865.69 |
| Phe-Tyr-Asp (and isomers) | ari | 0.01 | (-0.01– 0.02) | 0.35 | -2884.36 |
| | log | 0.30 | (-0.26– 0.85) | 0.31 | -2884.28 |
| 4-Methylhippuric acid | ari | 0.01 | (-0.01– 0.02) | 0.38 | -2729.32 |
| | log | 0.29 | (-0.26– 0.84) | 0.31 | -2729.11 |
| 3-Methylhippuric acid | ari | 0.01 | (-0.01– 0.02) | 0.38 | -2729.32 |
| | log | 0.29 | (-0.26– 0.84) | 0.31 | -2729.11 |
| Valerylglycine isomer 2 | ari | -0.01 | (-0.02– 0.01) | 0.32 | -2899.58 |
| | log | -0.28 | (-0.83– 0.28) | 0.34 | -2899.62 |
| Porphobilinogen | ari | -0.01 | (-0.02– 0.01) | 0.33 | -2807.25 |
| | log | -0.28 | (-0.83– 0.27) | 0.33 | -2807.26 |
| Pyruvic acid | ari | -0.01 | (-0.02– 0.01) | 0.33 | -2785.91 |
| | log | -0.27 | (-0.82– 0.28) | 0.35 | -2785.95 |
| Isovalerylalanine isomer | ari | -0.01 | (-0.02– 0.01) | 0.34 | -2459.47 |
| | log | -0.20 | (-0.75– 0.35) | 0.49 | -2459.63 |
| Glyceric acid | ari | -0.01 | (-0.02– 0.01) | 0.42 | -2630.90 |
| | log | -0.28 | (-0.83– 0.28) | 0.34 | -2630.73 |
| Pyrocatechol sulfate | ari | 0.01 | (-0.01– 0.02) | 0.35 | -3409.57 |
| | log | 0.26 | (-0.29– 0.82) | 0.37 | -3409.61 |
| N-Acetylisoleucine | ari | -0.01 | (-0.02– 0.01) | 0.35 | -2741.28 |
| | log | -0.25 | (-0.80– 0.30) | 0.39 | -2741.36 |
| 2-acetamido-4-methylphenyl acetate | ari | 0.01 | (-0.01– 0.02) | 0.36 | -2387.78 |
| | log | 0.25 | (-0.31– 0.80) | 0.40 | -2387.86 |
| Mevalonic acid isomer 1 | ari | -0.01 | (-0.02– 0.01) | 0.38 | -2426.41 |
| | log | 0.01 | (-0.54– 0.57) | 0.99 | -2426.91 |
| m-Hydroxyhippuric acid | ari | 0.00 | (-0.01– 0.02) | 0.52 | -3009.34 |
| | log | 0.25 | (-0.31– 0.80) | 0.40 | -3009.18 |
| Citric acid | ari | 0.01 | (-0.01– 0.02) | 0.48 | -3660.75 |
| | log | 0.24 | (-0.31– 0.79) | 0.41 | -3660.63 |
| 3,5-dihydroxybenzoic acid | ari | -0.01 | (-0.02– 0.01) | 0.46 | -2572.62 |
| | log | -0.23 | (-0.78– 0.32) | 0.44 | -2572.58 |
| 3,4-dihydroxybenzoic acid | ari | -0.01 | (-0.02– 0.01) | 0.46 | -2572.62 |
| | log | -0.23 | (-0.78– 0.32) | 0.44 | -2572.58 |
| Cinnamoylglycine | ari | 0.01 | (-0.01– 0.02) | 0.44 | -3174.17 |
| | log | 0.19 | (-0.36– 0.74) | 0.53 | -3174.27 |
| 3-Hydroxybenzyl alcohol | ari | -0.01 | (-0.02– 0.01) | 0.45 | -2558.06 |
| | log | -0.15 | (-0.71– 0.40) | 0.62 | -2558.23 |
| 3,4-Dihydroxybenzeneacetic acid | ari | -0.01 | (-0.02– 0.01) | 0.46 | -2579.44 |
| | log | -0.15 | (-0.70– 0.41) | 0.63 | -2579.62 |
| Salicylic acid | ari | 0.01 | (-0.01– 0.02) | 0.48 | -2435.10 |
| | log | 0.07 | (-0.49– 0.62) | 0.86 | -2435.25 |
| Glucuronic acid and/or isomers | ari | 0.01 | (-0.01– 0.02) | 0.49 | -2470.30 |
| | log | 0.15 | (-0.40– 0.70) | 0.62 | -2470.42 |
| Glycocholic acid isomer 2 | ari | -0.01 | (-0.02– 0.01) | 0.50 | -2222.84 |
| | log | 0.08 | (-0.48– 0.63) | 0.83 | -2223.29 |
| Gentisic acid | ari | -0.01 | (-0.02– 0.01) | 0.50 | -2785.85 |
| | log | -0.12 | (-0.67– 0.44) | 0.72 | -2786.03 |

| | | | | | |
|---|---|---|---|---|---|
| Azelaic acid | ari | -0.01 | (-0.02– 0.01) | 0.51 | -3004.31 |
| | log | -0.15 | (-0.70– 0.40) | 0.63 | -3004.39 |
| Nicotinuric acid isomer | ari | -0.01 | (-0.02– 0.01) | 0.52 | -2571.96 |
| | log | -0.14 | (-0.70– 0.41) | 0.63 | -2572.07 |
| Tetrahydrohippuric acid | ari | 0.01 | (-0.01– 0.02) | 0.52 | -2750.30 |
| | log | 0.14 | (-0.42– 0.69) | 0.66 | -2750.40 |
| Pyroglutamic acid | ari | 0.00 | (-0.01– 0.02) | 0.53 | -2757.24 |
| | log | 0.17 | (-0.38– 0.72) | 0.57 | -2757.28 |
| p-Hydroxyhippuric acid | ari | -0.00 | (-0.02– 0.01) | 0.60 | -3146.31 |
| | log | -0.18 | (-0.73– 0.37) | 0.54 | -3146.26 |
| 2-Isopropylmalic acid | ari | 0.00 | (-0.01– 0.02) | 0.56 | -2946.72 |
| | log | 0.10 | (-0.45– 0.65) | 0.74 | -2946.84 |
| Xanthosine | ari | -0.00 | (-0.02– 0.01) | 0.58 | -2689.75 |
| | log | -0.14 | (-0.70– 0.41) | 0.64 | -2689.78 |
| Taurine | ari | -0.00 | (-0.02– 0.01) | 0.8 | -2598.95 |
| | log | -0.16 | (-0.71– 0.39) | 0.6 | -2598.83 |
| 3-Methyl-2-oxovaleric acid | ari | -0.00 | (-0.02– 0.01) | 0.62 | -2892.88 |
| | log | -0.13 | (-0.68– 0.42) | 0.68 | -2892.93 |
| 3,3-Dimethylglutaric acid | ari | 0.00 | (-0.01– 0.02) | 0.83 | -2541.43 |
| | log | 0.13 | (-0.42– 0.68) | 0.68 | -2541.36 |
| Sulfosalicylic acid isomer | ari | -0.00 | (-0.02– 0.01) | 0.70 | -3077.88 |
| | log | -0.05 | (-0.60– 0.50) | 0.89 | -3077.96 |
| Ketoleucine | ari | -0.00 | (-0.02– 0.01) | 0.73 | -2732.25 |
| | log | -0.10 | (-0.66– 0.45) | 0.74 | -2732.27 |
| (2-methoxyethoxy)propanoic acid isomer | ari | -0.00 | (-0.02– 0.01) | 0.79 | -2485.21 |
| | log | 0.00 | (-0.55– 0.55) | 1.00 | -2485.31 |
| 2-Methylhippuric acid | ari | 0.00 | (-0.01– 0.02) | 0.89 | -2619.73 |
| | log | 0.09 | (-0.46– 0.64) | 0.79 | -2619.70 |
| 3-Hydroxyphenylacetic acid | ari | -0.00 | (-0.02– 0.01) | 0.96 | -2839.26 |
| | log | -0.08 | (-0.63– 0.47) | 0.81 | -2839.22 |
| Monoethyl phthalate | ari | 0.00 | (-0.01– 0.02) | 0.82 | -2512.30 |
| | log | 0.05 | (-0.51– 0.60) | 0.91 | -2512.33 |
| N-Acetylleucine | ari | -0.00 | (-0.02– 0.01) | 0.82 | -2541.55 |
| | log | -0.06 | (-0.61– 0.49) | 0.88 | -2541.57 |
| alpha-N-Phenylacetyl-glutamine | ari | -0.00 | (-0.02– 0.01) | 0.83 | -3396.04 |
| | log | -0.06 | (-0.62– 0.49) | 0.86 | -3396.05 |
| o-methylhippuric acid | ari | 0.00 | (-0.01– 0.02) | 0.83 | -2605.05 |
| | log | 0.02 | (-0.53– 0.57) | 0.97 | -2605.07 |
| m-methylhippuric acid | ari | 0.00 | (-0.01– 0.02) | 0.83 | -2605.05 |
| | log | 0.02 | (-0.53– 0.57) | 0.97 | -2605.07 |
| p-methylhippuric acid | ari | 0.00 | (-0.01– 0.02) | 0.83 | -2605.05 |
| | log | 0.02 | (-0.53– 0.57) | 0.97 | -2605.07 |
| p-Hydroxyphenylacetic acid | ari | -0.00 | (-0.02– 0.01) | 0.84 | -2650.88 |
| | log | -0.03 | (-0.58– 0.52) | 0.96 | -2650.91 |
| Chenodeoxycholic acid isomer | ari | 0.00 | (-0.01– 0.02) | 0.87 | -2384.89 |
| | log | 0.07 | (-0.48– 0.62) | 0.84 | -2384.88 |
| 9-Methylxanthine | ari | -0.00 | (-0.02– 0.01) | 0.84 | -3180.67 |
| | log | -0.06 | (-0.61– 0.50) | 0.88 | -3180.68 |
| Hydroxybenzyl alcohol isomer | ari | 0.00 | (-0.01– 0.02) | 0.89 | -2635.57 |
| | log | 0.07 | (-0.48– 0.62) | 0.85 | -2635.56 |

| | | | | | |
|---|---|---|---|---|---|
| 6-(2-hydroxyethoxy)-6-oxohexanoic acid | ari | -0.00 | (-0.02– 0.01) | 0.86 | -2346.24 |
| | log | -0.03 | (-0.58– 0.52) | 0.95 | -2346.26 |
| Malic acid | ari | 0.00 | (-0.01– 0.02) | 0.95 | -2555.95 |
| | log | 0.06 | (-0.49– 0.61) | 0.87 | -2555.94 |
| Hexanoylglycine | ari | -0.00 | (-0.02– 0.01) | 0.87 | -2710.24 |
| | log | -0.02 | (-0.57– 0.53) | 0.98 | -2710.26 |
| Phenol sulfate | ari | -0.00 | (-0.02– 0.01) | 0.98 | -3421.62 |
| | log | -0.05 | (-0.60– 0.50) | 0.89 | -3421.60 |
| Asp-Leu/Ile isomer 1 | ari | -0.00 | (-0.01– 0.01) | 0.99 | -2629.28 |
| | log | 0.04 | (-0.51– 0.59) | 0.92 | -2629.29 |
| Methyl (hydroxymethyl)pyrrolidine-carboxylate | ari | 0.00 | (-0.01– 0.02) | 0.96 | -2780.32 |
| | log | 0.02 | (-0.53– 0.57) | 0.98 | -2780.31 |
| Methyl (hydroxy)piperidine-carboxylate | ari | 0.00 | (-0.01– 0.02) | 0.96 | -2780.32 |
| | log | 0.02 | (-0.53– 0.57) | 0.98 | -2780.31 |
| Suberic acid | ari | -0.00 | (-0.01– 0.01) | 0.98 | -2811.84 |
| | log | 0.03 | (-0.52– 0.58) | 0.96 | -2811.83 |

Table 1: min p-Values of two models

Under a linear model (i.e., for the metameter "ari") investigate the association between age and all metabolites simultaneously

```
### linear effects only
make_str <- function(a)
  as.mlt(BoxCox(as.formula(paste(a, " ~ ageari")), data = dd))
res <- lapply(analyt, make_str)
names(res) <- analyt
### construct mmm / glht
a1 <- do.call("mmm", res)
a2 <- rep("ageari = 0", length(analyt))
names(a2) <- analyt
a2 <- as.list(a2)
a2 <- do.call("mlf", a2)
m <- glht(a1, a2)
### compute confidence internals
ci <- confint(m)
```

```r
ci <- ci$confint
ci.o <- ci[order(ci[,1]),]
ID <- which((ci.o[, 3] <= 0 | 0 <= ci.o[, 2]))
w.n<-rownames(ci.o[ID,])
w.n<-sub(": ageari","",w.n)
par(mar=c(5, 4, 4, 8) + 0.05)
plot(ci.o[,1], 1:nrow(ci.o), pch = 20, main="Metabolites vs. Age",
     xlab="Effects",ylab="Metabolites i=1,...,120")
colors <- rep("orange", nrow(ci.o))
colors[ID] <- "blue"
segments(ci.o[,2], 1:nrow(ci.o), ci.o[,3], 1:nrow(ci.o),col=colors)
abline(v = 0)
axis(4,at=which(colors=="blue"), labels=w.n,las=2,cex.axis=0.25,tck=-0.01)
```

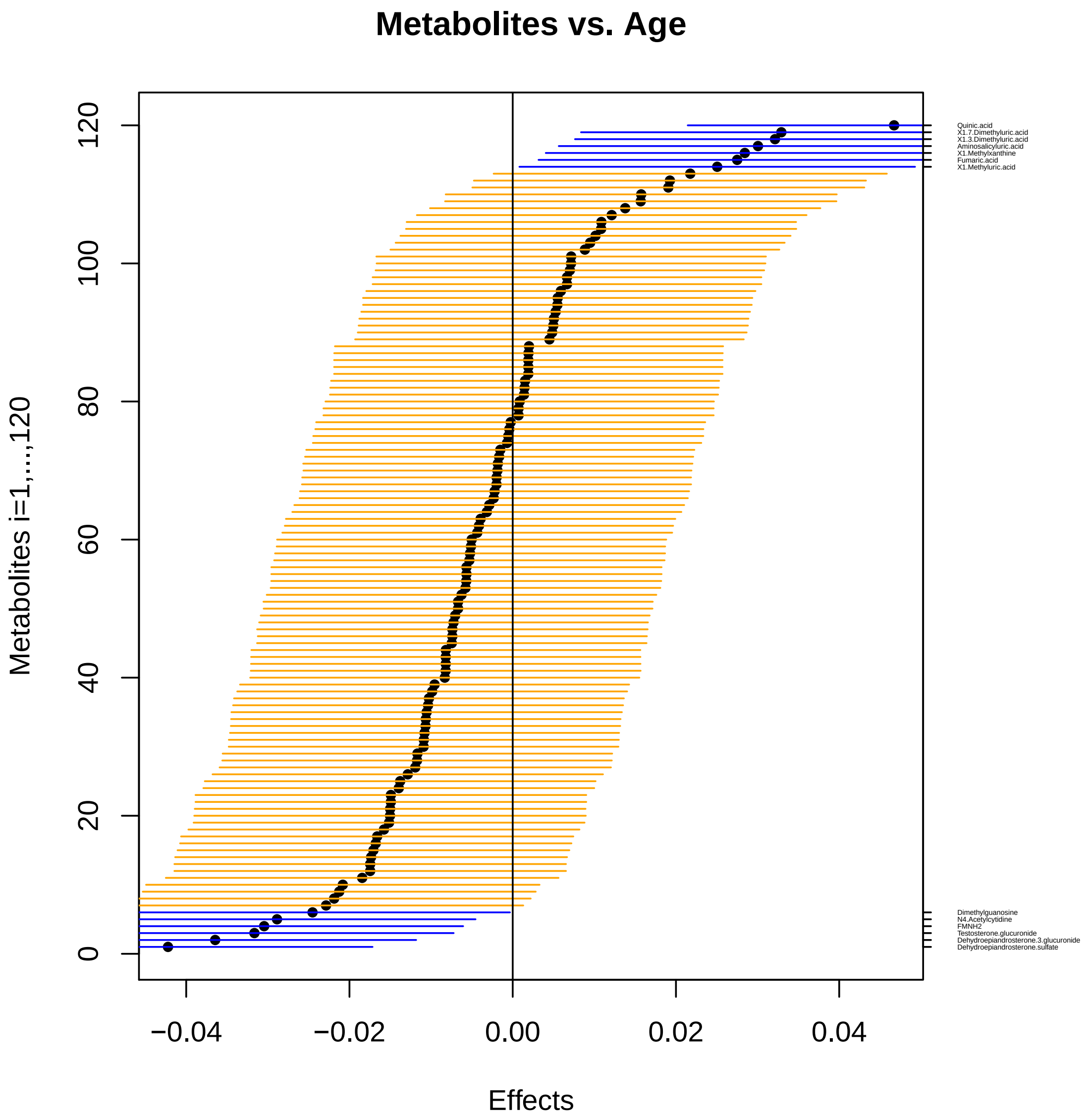


Figure 3: Impact of age on 120 urine metabolites for 183 adults under a linear model, visualized by confidence intervals, ordered by increasing effects, given as horizontal lines around black circles. The blue lines are the metabolites that show association with age.

```
## R version 3.6.0 (2019-04-26)
## Platform: x86_64-w64-mingw32/x64 (64-bit)
## Running under: Windows 10 x64 (build 17763)
##
## Matrix products: default
##
## locale:
## [1] LC_COLLATE=German_Germany.1252  LC_CTYPE=German_Germany.1252
## [3] LC_MONETARY=German_Germany.1252 LC_NUMERIC=C
## [5] LC_TIME=German_Germany.1252
##
## attached base packages:
## [1] stats     graphics  grDevices utils     datasets  methods   base
##
## other attached packages:
##  [1] gdata_2.18.0      data.table_1.12.2 tibble_2.1.3
##  [4] lattice_0.20-38   ggplot2_3.2.1     xtable_1.8-4
##  [7] tukeytrend_0.6    multcomp_1.4-10   TH.data_1.0-10
## [10] MASS_7.3-51.4     survival_2.44-1.1 mvtnorm_1.0-11
## [13] tram_0.2-6        mlt_1.0-5         basefun_1.0-5
## [16] variables_1.0-2   knitr_1.24
##
## loaded via a namespace (and not attached):
##  [1] gtools_3.8.1        tidyselect_0.2.5    zoo_1.8-6
##  [4] xfun_0.8            purrr_0.3.2         splines_3.6.0
##  [7] colorspace_1.4-1    mgcv_1.8-28         rlang_0.4.0
## [10] orthopolynom_1.0-5  nloptr_1.2.1        pillar_1.4.2
## [13] withr_2.1.2         glue_1.3.1          stringr_1.4.0
## [16] munsell_0.5.0       gtable_0.3.0        codetools_0.2-16
## [19] evaluate_0.14       labeling_0.3        pbkrtest_0.4-7
## [22] parallel_3.6.0      highr_0.8           Rcpp_1.0.2
## [25] polynom_1.4-0       scales_1.0.0        alabama_2015.3-1
## [28] lme4_1.1-21         BB_2014.10-1        digest_0.6.20
## [31] stringi_1.4.3       dplyr_0.8.3         numDeriv_2016.8-1.1
## [34] grid_3.6.0          quadprog_1.5-7      tools_3.6.0
## [37] sandwich_2.5-1      magrittr_1.5        lazyeval_0.2.2
## [40] coneproj_1.14       Formula_1.2-3       pkgconfig_2.0.2
## [43] crayon_1.3.4        Matrix_1.2-17       assertthat_0.2.1
## [46] minqa_1.2.4         R6_2.4.0            boot_1.3-22
## [49] nlme_3.1-139        compiler_3.6.0
```
