## Additional file 2: Rnw-Code that demonstrates the application of the workflow on a large-scale study (impact of age on the urinary metabolome). for "A new workflow combining R packages for statistical analysis of metabolites": re_analyse_thevenot.Rnw

```
\documentclass[11pt]{article}
\usepackage[a4paper,left=3cm,right=2cm,top=2.5cm,bottom=2.5cm]{geometry}
%% Load required LaTeX packages
%\usepackage{mathptmx}       % selects Times Roman as basic font
\usepackage{helvet}         % selects Helvetica as sans-serif font
%\usepackage{courier}        % selects Courier as typewriter font
\usepackage{graphicx} % standard LaTeX graphics tool
\usepackage{url}
\usepackage{enumerate}
\usepackage{amssymb, amsmath}
\allowdisplaybreaks
\usepackage{subfigure}
\usepackage[T1]{fontenc}
\usepackage{longtable}
\usepackage{float}
%% Set margins on highlighted code output boxes
\setlength\parindent{0pt}
\setlength\fboxsep{7mm}

%% include packages
<<prolog_packages, include=FALSE>>=
library("knitr")
library("tram")
library("tukeytrend")
library("multcomp")
library("survival")
library("xtable")
library("ggplot2")
library("lattice")
set.seed(01022019)
bread.mlt_fit <- function(x) vcov(x) * nrow(x$data) # sandwich estimator
bread.mlt <- function(x) vcov(x) * nrow(x$data) # sandwich estimator
@

%% Set knitr global options
 <<GlobalOpts, echo=FALSE, cache=FALSE,include=FALSE>>=
  opts_chunk$set(fig.align='center')
  opts_chunk$set(echo=TRUE)
  opts_chunk$set(tidy=FALSE)
  opts_chunk$set(cache=TRUE)
  opts_chunk$set(eval=TRUE)
@

\begin{document}
\begin{flushleft}
\textbf{Supplementary file of \\
''A new workflow combining R packages for statistical analysis of metabolites''}\\[0.6cm]
\begin{Large}
\textbf{Application of the workflow to a real, available dataset: Urinary metabolome variation with age}\\[0.6cm]
\end{Large}
\end{flushleft}
\tableofcontents


\section{Aim}
The aim of this report is to re-analyse large-scale data that are available in literature using the presented workflow. Thus, we demonstrate how the workflow works in practice, which assumptions are required and how the results can be represented in form of, for instance, confidence intervals, plots, lists of p-values, etc....The code is completely reproducible.\\
To this end, we considered a cohort consisting of 120 analytes measured by LC-HRMS for 183 adults (compare the section below). In particular, we focussed on the association between the analytes and age as continuous covariate of interest.

\section{Data}
Consider the article:\\
Th\'{e}venot et al. (2015) Analysis of the Human Adult Urinary Metabolome Variations with age, Body Mass Index, and Gender by Implementing a Comprehensive Workflow for Univariate and OPLS Statistical Analyses. \textit{J Proteome Res}, \textbf{14(8)}, 3322-3335.\\
DOI: \url{10.1021/acs.jproteome.5b00354 }\\
\url{https://pubs.acs.org/doi/10.1021/acs.jproteome.5b00354}
and the data available together with the article
\begin{itemize}
\item \texttt{sacurine-neg\_sampleMetadata.csv}, downloaded from\\
\url{https://workflow4metabolomics.org/dataset_sacurine} (October 31th, 2019)
\item \texttt{m\_sacurine\_v2\_maf.csv}, downloaded from \\
\url{https://www.ebi.ac.uk/metabolights/MTBLS404} (October 31th, 2019)
\end{itemize}

<<pfade,eval=TRUE,warning=FALSE,cache = TRUE, echo = FALSE>>=
#### metabolites table 
tf <- tempfile(fileext = "zip")
sysinf <- Sys.info()
os <- sysinf['sysname']
if (os == 'Windows') {download.file("https://www.ebi.ac.uk/metabolights/MTBLS404/files/m_sacurine_v2_maf.tsv",destfile = tf,method = "curl")}else{download.file("https://www.ebi.ac.uk/metabolights/MTBLS404/files/m_sacurine_v2_maf.tsv",destfile = tf)}
unzip(tf)
daten.1 <- read.table("m_sacurine_v2_maf.tsv")
# remove QC samples/blanks
daten.1<-daten.1[,-c(c(1:4),c(6:45))]
library("data.table")
daten.1.t<-transpose(daten.1)
colnames(daten.1.t)<-daten.1.t[1,]
daten.1.t<-daten.1.t[-1,]
colnames(daten.1.t)[1]<-"proband"
make.num.from.chr<- function(x) as.numeric((x))
daten.1.t[ ,2:dim(daten.1.t)[2]]<- 
lapply(daten.1.t[ ,2:dim(daten.1.t)[2]], make.num.from.chr)
daten.2 <- read.table("sacurine-neg_sampleMetadata.tsv",sep="\t",head=T)
daten.2<-daten.2[!is.na(daten.2$age),]
colnames(daten.2)[1]<-"proband"
daten.2<-daten.2[,-c(2:8)]
#### merge 
data.thevenot<-merge(daten.1.t,daten.2, by.x="proband",by.y="proband", all=FALSE)
# last three columns: age, BMI, gender
data.thevenot<-data.thevenot[,c(c(1,122,123,124),c(2:121))]
p<-dim(data.thevenot)[2]
name.analyt<-colnames(data.thevenot[,5:p])
@

\section{A little descriptive statistics}
%%% side-by-side; caption
Draw gender-specific age and BMI distributions
\begin{figure}
<<descr-age,echo=FALSE,fig.width=3,fig.height=3>>=
data.thevenot$gender<- factor(data.thevenot$gender, 
                              levels = c("M", "F"), labels = c("Male", "Female"))
ggplot(aes(y = age, x = gender), data = data.thevenot) +
geom_boxplot(outlier.size=1)+
xlab("Sex") +
ylab("Age")+
geom_jitter(position=position_jitter(width=0.2,height=0),
alpha=1,
size=1)
@
\caption{Gender-specific age distribution.}
\end{figure}
\begin{figure}
<<descr-bmi,echo=FALSE,fig.width=3,fig.height=3>>=
ggplot(aes(y = bmi, x = gender), data = data.thevenot) +
geom_boxplot(outlier.size=1)+
xlab("Sex") +
ylab("BMI")+
geom_jitter(position=position_jitter(width=0.2,height=0),
alpha=1,
size=1)
@
\caption{Gender-specific BMI distribution.}
\end{figure}

\newpage
\section{Re-analysation of data}
Is there any association between each metabolite and age or between each metabolite and log(age), respectively? \\ 
Run for this the workflow on the data 
<<analysis,echo=TRUE,warning=FALSE, cache = TRUE>>=
analyt <- make.names(analyt_orig <- colnames(data.thevenot)[-(1:4)])
colnames(data.thevenot)[-(1:4)] <- analyt

dd <- dosescalett(data = data.thevenot, dose = "age", 
                  scaling = c("ari","log"))$data

make_str <- function(a) {
  cm_ari <- as.mlt(BoxCox(as.formula(paste(a, " ~ ageari")), data = dd))
  cm_log <- as.mlt(BoxCox(as.formula(paste(a, " ~ agelog")), data = dd))
  cm_mmm <- glht(mmm(ari = cm_ari, log = cm_log),
                 mlf(ari = "ageari=0", log = "agelog=0"))
  list(cf = confint(cm_mmm)$confint, 
       pvals = summary(cm_mmm)$test$pvalue,
       logLik = c(ageari = logLik(cm_ari),
                  agelog = logLik(cm_log)))
}
res <- lapply(analyt, make_str)
@

<<table, results = "asis", echo = FALSE>>=
names(res) <- analyt_orig
or <- order(sapply(res, function(x) min(x$pval)))
res <- res[or]
rown <- rep(names(res), each = 2)
rown[duplicated(rown)] <- ""
col1 <- rep(c("ari", "log"), length(res))
xCI <- do.call("rbind", lapply(res, function(x) x$cf))
xCI <- formatC(xCI, digits = 2, width = 5, format = "f")
xCI <- cbind(xCI[,1], paste("(", xCI[,2], "--", xCI[,3], ")", sep = ""))
X <- cbind(xCI, do.call("c", lapply(res, function(x) format.pval(x$pvals, digits = 2, eps = 0.001))), 
                do.call("c", lapply(res, function(x) formatC(x$logLik, digits = 2, format = "f"))))
X <- cbind(rown, col1, X)
colnames(X) <- c("Analyte", "Age", "Effect", "Confint", "p-Value", "logLik")
library("xtable")
x.big.2 <- xtable(X,digits=c(5),
                  caption="min p-Values of two models")
print(x.big.2, hline.after=c(-1,0, cumsum(rep(2, length(analyt)))), include.rownames=FALSE,
       include.colnames=TRUE, floating = FALSE,
       tabular.environment = "longtable")
@

Under a linear model (i.e., for the metameter "ari") investigate the association between age and all metabolites simultaneously  

<<simultan_CI, cache = TRUE>>=
### linear effects only
make_str <- function(a)
  as.mlt(BoxCox(as.formula(paste(a, " ~ ageari")), data = dd))
res <- lapply(analyt, make_str)
names(res) <- analyt
### construct mmm / glht
a1 <- do.call("mmm", res)
a2 <- rep("ageari = 0", length(analyt))
names(a2) <- analyt
a2 <- as.list(a2)
a2 <- do.call("mlf", a2)
m <- glht(a1, a2)
### compute confidence internals
ci <- confint(m)
@

\begin{figure}
<<ci-plot>>=
ci <- ci$confint
ci.o <- ci[order(ci[,1]),]
ID <- which((ci.o[, 3] <= 0 | 0 <= ci.o[, 2]))
w.n<-rownames(ci.o[ID,]) 
w.n<-sub(": ageari","",w.n)
par(mar=c(5, 4, 4, 8) + 0.05)
plot(ci.o[,1], 1:nrow(ci.o), pch = 20, main="Metabolites vs. Age",
     xlab="Effects",ylab="Metabolites i=1,...,120")
colors <- rep("orange", nrow(ci.o))
colors[ID] <- "blue"
segments(ci.o[,2], 1:nrow(ci.o), ci.o[,3], 1:nrow(ci.o),col=colors)
abline(v = 0)
axis(4,at=which(colors=="blue"), labels=w.n,las=2,cex.axis=0.25,tck=-0.01)
@
\caption{Impact of age on 120 urine metabolites for 183 adults under a linear model, visualized by confidence intervals, ordered by increasing effects, given as horizontal lines around black circles. The blue lines are the metabolites that show association with age.}
\end{figure}
\newpage
<<sessionInfo, echo = FALSE>>=
sessionInfo()
@

\end{document}
```
