## Additional file 1: R-Code that demonstrates the workflow using the R packages survival, tram, mlt and multcomp, by means of the example data. for "A new workflow combining R packages for statistical analysis of metabolites": workflow_code.R

```
dd <-
structure(list(Age = c(66.3682409308693, 23.6448751642576, 
50.9869952087611, 23.3683749452475, 57.3100778852962, 64.8665532832787, 
69.249843554443, 30.1594983661574, 55.8419308357349, 64.8090596092421, 
55.066392881588, 61.1403338337896, 70.0444992866232, 58.1896055684455, 
23.0444900752909, 65.3401916455801, 21.1951219512195, 40.9815527019099, 
52.6584871222876, 64.3484120967063, 61.0554623333039, 54.1649659017373, 
49.5653854412712, 57.3401935331603, 62.6064339493498, 67.809070348768, 
70.7843942505133, 72.4086408640864, 75.9514311111111, 23.1101984941821, 
70.3730382138588, 59.6584982720704, 33.2854497141477, 32.238924838228, 
55.9268011527378, 49.0226140283634, 29.5653978627572, 26.7405886379192, 
33.6831195244055, 22.7241615331964, 73.0554589514965, 48.3018773047268, 
52.546238085581, 64.1567685957375, 68.5927722152691, 60.9979687362007, 
59.6995646514968, 61.5544304897658, 71.2005475701574, 64.5818227070975, 
48.1759414459716, 24.0144546649146, 58.4688631090487, 51.4743326488706, 
68.5181526008809, 67.5263518138262, 29.7762077188024, 48.5592244943569, 
65.0390343053885, 33.9240456821026, 59.2197125256674, 18.8720054757016, 
58.4935034802784, 53.4195903467856, 71.9514326432643, 26.4524437233827, 
23.5345653661875, 28.9158605584961, 23.6851471594798, 64.258065874821, 
60.7926344608319, 24.5537669732808, 25.340212698747, 62.4414845074095, 
22.8747433264887, 50.1321129482499, 26.4716081930643, 31.7789945246391, 
45.7625094658356, 62.3757768023989, 62.8090349075975, 21.6714783474365, 
60.7597809767729, 65.3046003235575, 58.6988399071926, 60.052735514564, 
61.6523006270423, 20.7207665232695, 24.471638195357, 50.4661262615418, 
69.8254733351328, 52.7953761914419, 62.0006953196297, 45.9541562299761, 
80.4332792537011, 60.8282257352292, 71.9185793579358, 74.9678302532512, 
62.9377138945927, 63.1047227926078, 67.4688569472964, 70.9760438056126, 
37.3402017291066, 21.4031856645097, 66.6358695652174, 61.5345756424976, 
58.2470997679814, 61.9952196775455, 71.6550308008214, 60.5701718953368, 
39.1457905544148, 73.7043175848163, 70.082828828134, 77.4250614250614, 
58.4880278422274, 61.1047425593924, 64.2005728245304, 57.1430729289592, 
74.3182813754837, 66.4223193854201, 61.4716064647178, 73.7097931850975, 
43.6139630390144, 26.3237679983776, 25.0773928609034, 62.7590717482943, 
70.1211583696448, 68.8220449946435, 63.378507871321, 60.8829815419942, 
60.4332839639154, 55.2224503764545, 72.4798229822982, 66.2881660673423, 
61.7891901439548, 53.6413506388157, 59.62765229295, 70.0171210426869, 
76.4168533333333, 49.2224716640201, 55.9459654178674, 51.2662559890486, 
64.258065874821, 64.1157021312442, 65.1704484174721, 48.2279584311096, 
56.3347262247839, 67.4633812457221, 69.9733158523889, 62.984257357974, 
53.981532181791, 51.5071868583162, 66.4004168028768, 69.3922090112641, 
63.5674195756331, 64.4853003116839, 58.9397679814385, 61.4332773999823, 
77.0034398034398, 54.2635273034994, 25.3929223281282, 25.7926384100588, 
79.5188102078756, 20.129335071708, 66.4585900068446, 62.2450376454483, 
65.4859839816934, 48.7105075836391, 51.6447463580948, 56.272456927071, 
57.8555916473318, 73.9733153245236, 26.3408624229979, 22.7104722792608, 
27.206665407855, 79.025325119781, 69.0171307884856, 41.9130905386477, 
54.2614647501711, 55.2696575243768, 47.74604682349, 24.0882707260979, 
36.5818443804035, 24.0444639141387, 55.0280629705681, 60.365539168065, 
45.9651074736413, 22.6611909650924, 20.2122946739672, 45.3627890720569, 
47.0636550308008, 43.732355804332, 23.6146095717884, 25.7324072196309, 
35.5735533520793, 27.4914086102719, 50.0362894567318, 23.9212572555032, 
32.6639826072953, 53.1239099574123, 23.2477275216296, 27.4777190332326, 
46.6584531143053, 41.2395899656182, 23.1266255989049, 42.0855723927162, 
57.2258932714617, 36.2826698238863, 50.9158110882957, 23.0828199863107, 
45.579076134444, 31.8309134655273, 42.0554565134344, 22.4093086926762, 
45.7652472767519, 31.970546751846, 23.3216515168109, 36.121133639189, 
34.4695414099932, 66.3490759753593, 23.9239951812507, 39.1430527036277, 
46.3928815879535, 52.3600689515311, 21.1239422598308, 25.830967349422, 
58.8446269678303, 25.0007370748657, 21.1814335490294, 43.0527036276523, 
46.252577619852, 32.4312746597955, 27.8117447129909, 24.6092450247447, 
61.9267741514928, 62.1622267611142, 50.1074726218596, 24.2525462709451, 
61.5626439528921, 24.6119827313889, 21.8528746577788, 35.2470910335387, 
47.8008046041236, 30.3874058863792, 50.0499785269487, 44.3792405665992, 
31.4804612959429, 38.8993839835729, 47.2306639288159, 32.806345116354, 
38.9650924024641, 30.5763175906913, 34.6255989048597, 31.611880859537, 
29.7789455091407, 40.3730217028381, 26.9541409993155, 41.5106328791545, 
48.8309697201993, 49.5790745114881, 46.5681040383299, 29.129403175762, 
32.2580270472082, 42.1375907296575, 25.1102453406339, 24.7543434768874, 
34.6475017111567, 30.1540227854809, 48.4551600826954, 28.836466508487, 
30.5516769336071, 43.9048430275006, 27.5023602719033, 46.735112936345, 
34.2991239048811, 29.7296652830522, 29.3812739551013, 23.4524298425736, 
47.1238877481177, 29.7679943477877, 32.104704223015, 25.9185763536808, 
22.8172484599589, 34.0116551939925, 34.1786608260325, 29.2252235809454, 
24.5181250684481, 29.4853075378719, 38.3812565812566, 23.9897053991896, 
27.5406910876133), GUDCA = c(221.6, 940, 72.3, 823.3, 111.3, 
222.4, 311.2, 114, 195.8, 80, 150.8, 25, 38.9, 144.5, 231.4, 
63.3, 196.8, 44.2, 27.2, 76.8, 169.5, 339.4, 25, 41.1, 395, 268.2, 
59, 614.7, 66.7, 151.2, 101, 72.2, 52.5, 506.2, 311.4, 150.5, 
226.5, 162.2, 61.9, 51.2, 68.4, 90, 39.1, 54.8, 87.7, 176, 25, 
367.8, 28.1, 53.2, 41.9, 45.3, 85.7, 98, 61.5, 35, 66.7, 51.2, 
107.3, 140.1, 141.9, 61.7, 369.3, 74.4, 38.1, 91.5, 119.9, 263, 
53, 374.8, 139.4, 900.9, 373, 91.2, 43.5, 74.6, 143.7, 64.7, 
49.3, 25, 51.4, 52.6, 164.9, 64.5, 120, 25, 45.8, 250.7, 88.5, 
44.8, 70.2, 162.3, 50.1, 26.9, 77.7, 26.4, 25, 33.3, 55.1, 126.6, 
43.3, 61.2, 299.2, 867.4, 290.8, 32.3, 57.6, 25, 212.3, 169.7, 
92.2, 25, 53.6, 28.5, 25, 149, 37.4, 282.5, 43.6, 25, 70.1, 26, 
84.3, 351.2, 31.4, 178.8, 35.6, 47.8, 322.1, 85.1, 96.1, 110.4, 
31.7, 376.1, 491.7, 25, 25, 59.5, 43.7, 25, 205.4, 25, 41.8, 
25, 63.1, 221.8, 25, 114.9, 83.5, 110.1, 310.8, 25, 135.5, 248.8, 
25, 31, 47.5, 45.6, 142.6, 45.4, 120.9, 95.8, 111.3, 66.5, 154.8, 
75.2, 25, 45.6, 242.7, 54.6, 76.2, 25, 52.5, 37.3, 42.5, 38.2, 
61.1, 152.5, 62.5, 64.9, 58.3, 210, 86, 32.4, 49.3, 25, 69.6, 
59.9, 593.7, 25, 81.8, 118.8, 69.7, 41, 913.8, 144.4, 53.8, 25, 
164.5, 97.1, 99.9, 120, 153.2, 70.1, 447.1, 43.6, 226.7, 25, 
256, 243.4, 240.5, 260.1, 142, 695, 47.9, 25, 243.3, 319.7, 104.4, 
31, 46.9, 213, 25, 132, 422.6, 310, 76.4, 331.7, 90.6, 25, 701.6, 
496.8, 426.6, 124.6, 27, 25, 38, 28.4, 61.3, 194.5, 488.8, 102.8, 
149.3, 37.9, 25, 141.1, 61.9, 184.6, 25, 28.8, 25, 355.1, 178, 
37.1, 35.4, 30.1, 597.1, 82.4, 55.7, 25, 58.8, 25, 177.3, 40.8, 
60.9, 80.6, 105.6, 284.5, 25, 186.2, 60.1, 25, 109.1, 83.8, 241, 
45.4, 154.6, 192.4, 133.5, 36.1, 50.5, 209.5, 66.5, 102.2, 308.2, 
134.7, 219.6, 462.9, 220.2, 254.2, 438.3), C10.0 = c(0.46, 0.54, 
0.46, 0.46, 1.52, 0.53, 2.1, 0.46, 0.46, 0.46, 0.46, 0.46, 0.46, 
0.65, 0.8, 0.46, 0.46, 0.48, 0.46, 0.61, 0.88, 0.43, 0.46, 0.46, 
0.43, 0.46, 0.51, 0.46, 0.46, 0.46, 0.67, 0.48, 0.46, 0.7, 0.46, 
0.46, 0.46, 0.74, 0.46, 0.46, 0.67, 0.82, 0.73, 0.51, 1.09, 0.46, 
0.46, 0.51, 1.57, 0.46, 0.97, 0.46, 0.46, 0.46, 0.46, 0.46, 0.46, 
1.97, 0.81, 0.83, 0.46, 0.46, 0.88, 0.46, 0.46, 0.57, 0.46, 3.66, 
0.46, 0.55, 0.74, 0.46, 0.46, 0.46, 0.51, 1.71, 0.52, 0.46, 0.46, 
0.46, 0.54, 0.46, 0.46, 1.71, 1.06, 0.57, 0.59, 3.79, 0.46, 0.46, 
0.46, 0.64, 0.46, 0.64, 0.57, 0.43, 0.46, 0.46, 0.67, 0.59, 0.63, 
0.48, 1.32, 0.67, 0.46, 0.86, 0.46, 0.46, 0.46, 0.53, 0.46, 0.46, 
0.79, 0.46, 0.66, 0.46, 0.46, 0.46, 0.46, 0.46, 0.46, 0.46, 0.46, 
0.67, 0.46, 0.46, 0.46, 0.46, 0.46, 0.46, 0.46, 0.46, 0.46, 1.1, 
0.46, 0.46, 0.46, 0.5, 0.46, 0.46, 0.46, 0.46, 0.46, 0.46, 0.51, 
0.68, 0.46, 0.46, 0.46, 0.78, 0.46, 0.46, 2.43, 1.61, 0.46, 0.46, 
0.46, 0.46, 0.62, 0.46, 0.54, 0.46, 0.46, 0.46, 1.29, 0.46, 0.46, 
0.46, 0.8, 0.46, 0.52, 0.46, 0.46, 0.46, 1.14, 0.59, 0.46, 0.46, 
3.6, 0.46, 0.46, 0.46, 0.46, 0.46, 0.49, 0.46, 0.46, 0.61, 0.46, 
0.63, 0.46, 0.46, 0.46, 1.03, 0.46, 0.46, 0.46, 0.46, 0.02, 0.68, 
0.46, 0.46, 0.43, 0.46, 0.46, 0.46, 0.46, 0.57, 0.46, 0.46, 0.47, 
0.46, 0.96, 0.79, 0.46, 0.46, 0.46, 0.47, 0.46, 0.46, 0.46, 0.46, 
0.46, 0.46, 0.65, 1.56, 1.04, 0.46, 0.46, 0.66, 0.46, 0.46, 0.46, 
0.46, 0.46, 0.46, 0.46, 0.46, 0.46, 0.45, 0.46, 0.46, 0.77, 0.46, 
0.46, 0.46, 0.46, 0.46, 0.55, 0.46, 0.46, 0.61, 0.74, 0.46, 0.46, 
0.46, 0.46, 0.46, 0.46, 0.46, 0.73, 0.46, 0.46, 0.6, 0.46, 0.46, 
0.5, 0.46, 0.46, 0.46, 0.56, 0.46, 0.73, 0.46, 0.46, 0.46, 0.81, 
0.46, 0.46, 0.46, 0.7, 0.46, 0.46, 0.02, 0.46, 0.46, 0.46, 0.46, 
0.66, 0.55, 0.46)), .Names = c("Age", "GUDCA", "C10.0"), class = "data.frame", row.names = c(NA, 
-291L))

### attache required packages
library("tram")
library("tukeytrend")
library("multcomp")
library("survival")
library("xtable")
set.seed(290875)

### define bread part for sandwich estimation of vcov's
bread.mlt_fit <- function(x) vcov(x) * nrow(x$data) 
bread.mlt <- function(x) vcov(x) * nrow(x$data) 

### (I) endpoint GUDCA with left censoring at value 25
(dd$GUDCAsurv <- with(dd, Surv(GUDCA, event = GUDCA > 25, type = "left")))
## support of baseline transformation
suGUDCA <- quantile(survfit(GUDCAsurv ~ 1, data = dd), 
                    prob = c(0, .95))$quantile
## age metameters
TukeyMetam <- dosescalett(data = dd, dose = "Age", 
                          scaling = c("ari", "log"))
m_dat <- TukeyMetam$data # Tukeys dose metameters
## transformation models: h(GUDCA) ~ N(beta * Tukey(age), 1)
fma <- as.mlt(BoxCox(GUDCAsurv ~ Ageari, data = m_dat, 
                     support = suGUDCA))
fmA <- as.mlt(BoxCox(GUDCAsurv ~ Agelog, data = m_dat,
                     support = suGUDCA))
## joint inference for all two metameters
maxT <- glht(mmm(ari = fma, log = fmA),
             mlf(ari = "Ageari=0", log = "Agelog=0"))
## single-step adjusted p-values
summary(maxT)
## global test
(minpGUDCA <- min(summary(maxT)$test$pvalue))
library(ggplot2)

### (II) Endpoint C10.0 with no censoring but many ties. Treat data as
### interval-censored
dd$C10.0surv <- with(dd, Surv(time = C10.0 - 0.005, time2 = C10.0 + 0.005, 
                              type = "interval2"))
suC10.0 <- quantile(survfit(C10.0surv ~ 1, data = dd), 
                    prob = c(.01, .9))$quantile
m_dat$C10.0surv <- dd$C10.0surv

fma2 <- as.mlt(BoxCox(C10.0surv ~ Ageari, data = m_dat, 
                      support = suC10.0))
fmA2 <- as.mlt(BoxCox(C10.0surv ~ Agelog, data = m_dat,
                      support = suC10.0))

maxT2 <- glht(mmm(ari = fma2,log = fmA2),
             mlf(ari = "Ageari=0", log = "Agelog=0"))
## single-step adjusted p-values
summary(maxT2)
## global test
(minpC10.0 <- min(summary(maxT2)$test$pvalue))

####### bivariate Multiple endpoints test
maxTbiv <- glht(mmm(ari=fma, log=fmA, ari2=fma2, log2=fmA2),
           mlf(ari="Ageari=0", log="Agelog=0",
               ari2="Ageari=0", log2="Agelog=0"))
multEndpoint<-summary(maxTbiv)
CIBB<-confint(maxTbiv)

confint(maxTbiv)
```
